# VitriFlex: An Open-Source, Modular, and Customizable Robotic Platform for Cryo-EM Grid Preparation

**DOI:** 10.1101/2025.07.24.666634

**Authors:** Wyatt Peele, Kedar Sharma, Thomas B. Stanley, Randy K. Bledsoe, Robert M. Petrovich, Mario J. Borgnia, Venkata P. Dandey

## Abstract

Specimen preparation remains the principal bottleneck in cryo-electron microscopy (cryo-EM). Achieving uniformly thin ice without denaturation, preferred particle orientation, or air– water-interface damage is challenging. The demands of time-resolved experiments and sub-nanoliter sample volumes raise the bar even further. Commercial plungers are closed, high-cost platforms that limit experimental flexibility, constraining users to a few vendor-defined “custom” routines. By contrast, an open, modular and automated system would allow researchers to tailor experimental conditions by tweaking mixing, dispensing, and freezing parameters while lowering barriers to adoption.

We present VitriFlex, an open-source cryo-EM grid preparation platform built around an industrial-grade SCARA robot and 3D-printed components. Designed for modularity and ease of assembly, the system enables programmable control of grid handling, sample deposition, and blotting, supporting both standard and time-resolved workflows.

VitriFlex’s modularity allows flexible implementation of sample delivery strategies, including pre-mixing and on-grid mixing modes, combined with acoustic-assisted spray application. The system allows for spray-to-plunge delays as short as ∼130 ms, enabling initiation of biochemical reactions within sub-second timescales prior to vitrification. Protocols are easily modifiable, and new components can be integrated with minimal redesign.

We validated the platform using standard single-particle samples including apoferritin and dGTPase. Additionally, we demonstrate effective mixing capabilities by generating spike–ACE2 and α7–bungarotoxin complexes. In all cases, VitriFlex reproducibly yielded high-quality, collectable grids, allowing us to achieve high-resolution reconstructions. These results highlight VitriFlex as a flexible, accessible solution for both routine and time-resolved cryo-EM applications.

## Introduction

Cryo-electron microscopy (cryo-EM) has transformed structural biology by enabling near-atomic resolution imaging of complex macromolecular assemblies without the need for crystallization^1–4^. Preserving the native structure of these targets under the vacuum and radiation damage of an electron microscope requires immobilizing them in a thin, amorphous layer of ice and maintaining them at ultracool temperatures. This is achieved through vitrification, a rapid freezing process that prevents the formation of damaging ice crystals while preserving the native hydration and structural integrity of biological macromolecules. The most common method for vitrification is blot and plunge freezing. In this technique, an aqueous specimen is usually applied to a perforated film supported by a grid and excess solution is blotted away resulting in a layer thinner than 100 nm^5^. The grid is then rapidly plunged into a cryogen, usually liquid ethane or an ethane-propane mixture cooled by liquid nitrogen. The ultra-rapid cooling rate (10□–10□ K/s) transforms the water into a non-crystalline glass-like state, trapping macromolecules in their native conformation. Plunge freezing is the foundational first step in cryo-EM workflows and is critical for achieving high-resolution structural data.

Despite its central role, plunge freezing remains one of the least standardized and most variable steps in cryo-EM specimen preparation. A wide range of commercial and open-source devices has been developed to address this challenge, offering differing degrees of automation, environmental control, and flexibility. However, no single system fully resolves the competing demands of reproducibility, adaptability, cost, and ease of use. While many refinements to plunge freezing have been developed, such as pin printing^6^ and self-wicking grids^7^, true alternatives remain limited. High-pressure freezing, which is suitable for thick cellular specimens, is not applicable to single-particle cryo-EM because the resulting specimen is too thick for direct imaging^8^. Jet vitrification, as implemented in systems like VitroJet^9^, represents an advanced variant of plunge freezing rather than a distinct method.

Plunge-freezing systems range from fully manual to fully automated designs. Manual plungers are simple and inexpensive to construct and offer experimental flexibility but require considerable operator skill and attention to environmental parameters to achieve reproducible results. Commercially available systems such as the Leica EM GP and Vitrobot Mark IV offer automation and environmental control, thereby improving consistency but limiting user customization. More recent systems, including Chameleon^10^ and VitroJet, provide full automation with minimal specimen consumption. Chameleon uses picoliter inkjet dispensers to reduce specimen volumes but is prone to clogging and requires specialized self-wicking grids, introducing additional complexity and cost. VitroJet employs pin printing for small-volume specimen application but remains expensive and lacks modularity, which can limit accessibility and adaptation for novel protocols.

Open-source efforts have addressed some of these limitations by offering modularity and affordability. Simple robotic plungers employing solenoids or servo motors provide single-axis motion and enable automated plunging but lack versatility. Spotiton^11,12^, an open-source system with a 3-axis robotic arm, improves automation, grid handling and mixing on grid^13^ using two picoliter dispenser but remains technically demanding to build and maintain. Using microfluidic systems two samples are efficiently mixed and dispensed on to the grid for time-resolved studies^14–18^. Alternative approaches such as Back-It-Up (BIU)^19^ and Shake-It-Off^20^ use acoustic transducers to deliver specimens, eliminating clogging by avoiding nozzle-based delivery. However, the absence of precision robotics restricts these systems to relatively simple protocols and reduces their suitability for complex or reproducible workflows. Although rapid spray-to-plunge methods minimize particle exposure to the air-water interface^21–27^, further improvements in reproducibility, flexibility and reasonable cost to build are needed for widespread adoption.

To address these challenges, we have developed an automated specimen preparation platform that provides a balance between automation, modularity, and user control. Our system is based on a commercially available SCARA robotic arm with plug-and-play capability and a built-in scripting language, offering precise, reproducible operation and easy integration into existing workflows. The robotic platform is mounted on an optical breadboard, allowing straightforward incorporation of external components for customized applications. The system is controlled by open-source software, enabling researchers to easily adapt protocols as needed. This design offers hands-free grid handling, efficient storage, and an accessible price point compared to available commercial systems.

We demonstrate the flexibility and robustness of the system by performing a range of specimen preparation workflows, including conventional blot-and-plunge, pre-mixing on an acoustic transducer, on-grid mixing, and rapid spray-to-plunge freezing within ∼350 milliseconds. In these experiments, high-resolution cryo-EM structures were obtained (Table 1), highlighting the system’s performance across a range of specimen types and protocols. By combining adaptability with accessibility, our modular platform with precision robotics empowers researchers to tailor specimen preparation protocols to the unique requirements of their biological specimens, contributing to enhanced reproducibility and throughput in cryo-EM studies.

**Table 1.**
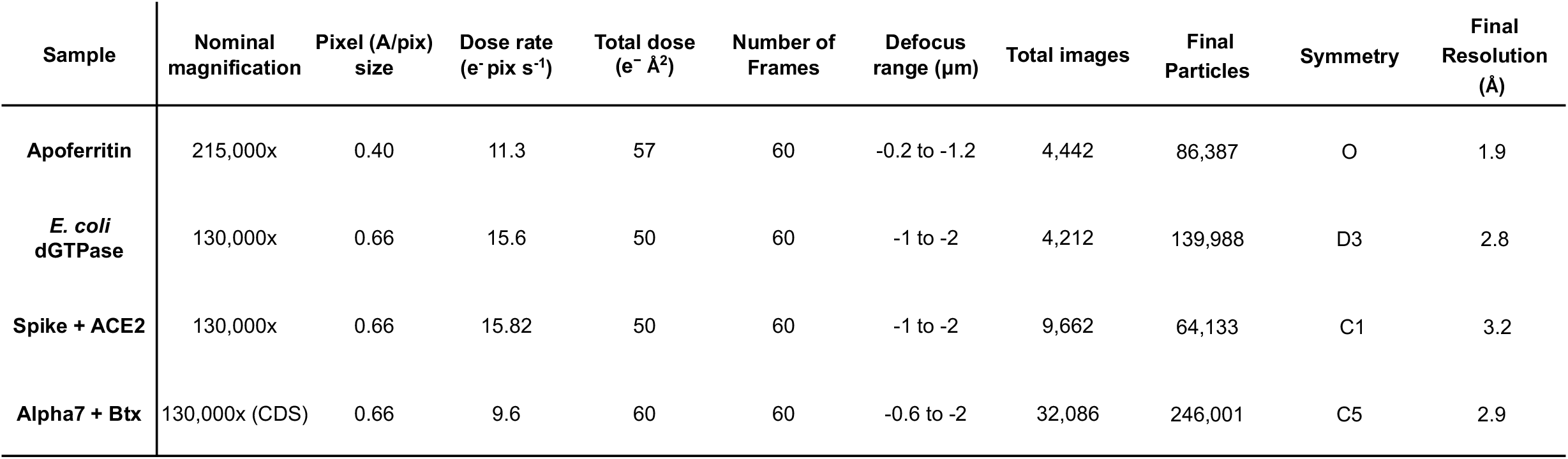
Collection parameters for cryo-EM structural determination.

| Sample | Nominal magnification | Pixel (Å/pix) size | Dose rate (e<sup>-</sup> pix s<sup>-1</sup>) | Total dose (e<sup>-</sup> Å<sup>2</sup>) | Number of Frames | Defocus range (μm) | Total images | Final Particles | Symmetry | Final Resolution (Å) |
| --- | --- | --- | --- | --- | --- | --- | --- | --- | --- | --- |
| **Apo ferritin** | 215,000x | 0.40 | 11.3 | 57 | 60 | -0.2 to -1.2 | 4,442 | 86,387 | O | 1.9 |
| ***E. coli* dGTPase** | 130,000x | 0.66 | 15.6 | 50 | 60 | -1 to -2 | 4,212 | 139,988 | D3 | 2.8 |
| **Spike + ACE2** | 130,000x | 0.66 | 15.82 | 50 | 60 | -1 to -2 | 9,662 | 64,133 | C1 | 3.2 |
| **Alpha7 + Btx** | 130,000x (CDS) | 0.66 | 9.6 | 60 | 60 | -0.6 to -2 | 32,086 | 246,001 | C5 | 2.9 |

## Materials and Methods

### System Overview and Design

All cryo-EM grids for this study were prepared on the VitriFlex platform (Fig. 1), an open-source, breadboard-mounted freezing system that combines a SCARA robot, a modular specimen-preparation chamber, and custom control software. The robot is an Epson T3 SCARA arm (repeatability ± 0.020 mm; four degrees of freedom; vertical-axis speed 1 m sβ^1^) connected to a Windows control PC via USB. The arm is fixed to the breadboard via a 3-D-printed spacer, and its digital and analog I/O lines are accessed through an Epson API to synchronize peripheral devices.

**Figure 1.**
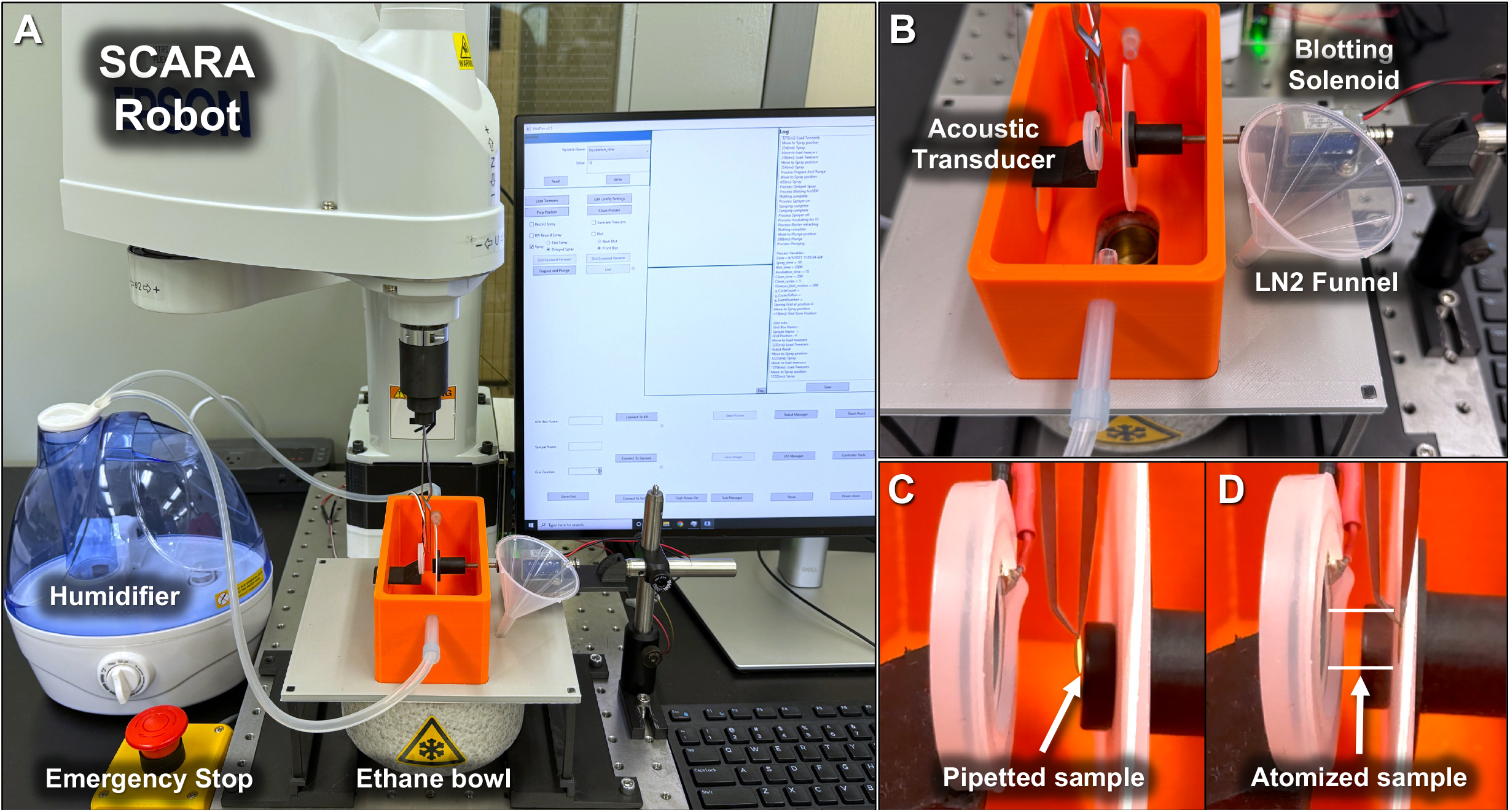
VitriFlex device setup. (A) The SCARA robot, controlled by a Windows PC, mounted to an optical breadboard along with a 3D printed specimen preparation chamber. (B) Close up of the preparation chamber surrounding the grid to hold humidity. (C) View of a grid placed in the “Prep” position with a droplet applied. (D) A sample being sprayed onto the grid during blotting.

The 3D-printed specimen preparation chamber integrates three core components: 1) a blotting device holds standard 47 mm filter paper discs (e.g., Whatman 40) and is actuated by a push-type solenoid. 2) a non-contact sample application system uses an ultrasonic piezoelectric transducer to deliver a fine aerosol spray onto the grid. 3) a 3D-printed cryogen holder accommodates the standard Vitrobot ethane cup, with an alternative configuration available for compatibility with the Nanosoft cryostat. Additionally, two lateral inlet ports support the connection of a humidifier, conditioning the environment around the grid during incubation.

System operation is managed through an open-source graphical user interface (GUI) developed in C#. Users can define spray, blot, and incubation timing, and train robotic positions using a jog-to-record function. Each run is automatically logged to a plain-text file, recording the date, time, grid ID, protein sample, box position, and all timing parameters. For enhanced safety, a large physical emergency-stop button mounted on the breadboard immediately cuts power to the robot.

Complete CAD designs (modeled using Autodesk Fusion 360), STL files, wiring diagrams, and source code are provided in the Supplementary Build Guide and on the project GitHub repository (**<u>GitHub URL</u>**). All printed parts were produced in a blend of nylon (specifically PA6) and micro-carbon fibers filament on a Markforged Mark Two printer; ABS and PETG are suitable alternatives.

### Sample Delivery and Blotting

Prior to freezing, all grids were rendered hydrophilic through surface treatment. Carbon grids were glow-discharged in air for 30 seconds using a PELCO easiGlow system. Gold grids were plasma treated in immersion mode with a gas mixture of oxygen and argon (1:9 sccm) at 38 W for 75 seconds using a PIE Scientific Tergeo EM.

The system supports multiple sample application modes. Specimens can be applied directly to the grid using a pipette or delivered via the integrated sprayer module. Blotting, spraying, and incubation durations are user-defined based on experimental requirements, and each function is executed sequentially once the protocol is initiated. Currently implemented protocols include “Spray” and “Blot.” Within the spray protocol, users can choose between a fast or a delayed option. In “fast spray”, the sample is applied immediately after blotting begins, with any incubation time appended after the blotting cycle. In contrast, “delayed spray” introduces incubation prior to blotting. All operations are controlled through a graphical user interface (GUI) and the robot’s I/O lines via the Epson API.

#### Blot and Plunge Protocol

All movements of the robotic arm and actions of other components are controlled from the GUI (Fig. S1). To initiate the blot-and-plunge freezing process, a pre-cleaned grid is gently secured using inverse tweezers. The robot’s arm is guided to the “Load Tweezers” position, an open, obstruction-free area selected to minimize any risk of damaging the grid and attach the tweezers to the robotic arm using an adapter and lock them in place. Once the tweezers are secured, the “Prep position” button moves the grid to the location where sample can be applied.

The blotting parameters including blot orientation and duration are configured at the beginning of the session. To begin, a small volume of sample (1-3µL) is applied to the application side of the grid using a pipette. With everything prepared, the “Prepare and Plunge” button triggers the blot and plunge sequence, rapidly freezing the sample into a thin, glass-like layer suitable for high-resolution analysis.

After vitrification in liquid ethane, the user can designate a storage slot and select “Store Grid”. This command prompts the robot to transfer the tweezers from ethane to liquid nitrogen, carefully maintaining a user-defined clearance to avoid damaging the grid, and then position the grid above the chosen storage location.

#### Spray-and-Plunge Mixing Protocol

The spray system is equipped with a piezoelectric transducer mounted approximately 0.5 cm from the grid, held in place by a removable silicone ring for easy cleaning. To prepare for each grid, 1–3 µL of the sample is applied onto the of the transducer—the side facing the grid. After spraying, the transducer surface is cleaned with 100% ethanol to ensure the droplet is placed centrally. This helps to avoid uneven spreading and ensures consistent spray delivery.

Alternatively, a small volume of buffer (1–3 µL) can be pre-applied to the grid before spraying to increase the number of collectable areas. The concentration of the sample sprayed onto the grid is adjusted accordingly. Once the buffer is in place, the sample is sprayed onto the wetted grid, allowed to incubate for the selected time, and then plunged (Fig. 2). This improves the spreading of liquid across the grid. This approach can be also used to mix two components on the grid. The first sample is pipetted directly onto the grid, whereas the second is applied onto the sprayer. Activation of the sprayer during the blotting cycle initiates the mixing of the two components.

**Figure 2.**
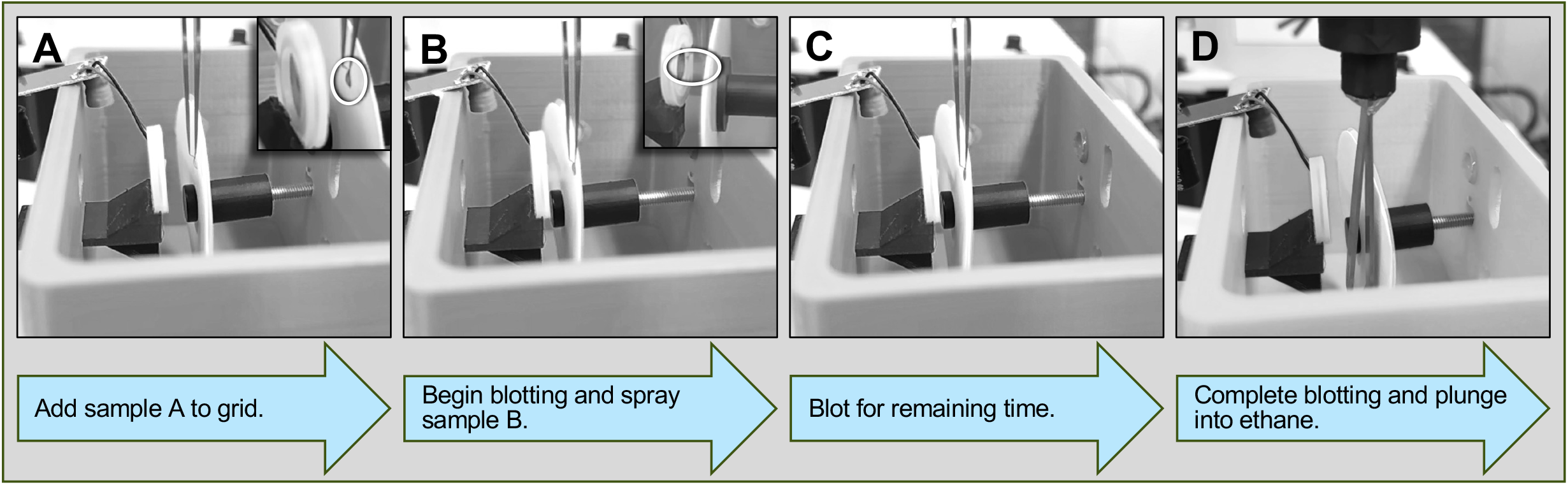
Time-resolved cryo-EM sample application and freezing workflow. (A) Sample A is applied to the application side of a pre-cleaned grid. (B) Blotting is initiated, and a pre-deposited Sample B is sprayed onto the grid. (C) Blotting continues for the remaining time, allowing sample incubation. (D) At the end of the blotting cycle, the blotting paper retracts, and the grid is plunged into liquid ethane.

### Preparation of Biological Samples

*Apoferritin* (400kDa; Sigma Aldrich, A3660, 2.5mg/ml) protein solution stored in 50% glycerol is exchanged into a cryo compatible buffer (50 mM Tris-Cl, pH 7.6; 150 mM NaCl) using Amicon Ultra-15 centrifugal filter units (100 kDa cutoff membrane).

*dGTPase* was purified following the method outlined previously.^28^

*SARS-CoV-2 S-protein* purification was done using a construct containing the prefusion Spike ectodomain (residues 1−1208) of 2019-nCoV S (GenBank:MN908947) with proline substitutions at residues 986 and 987, a “GSAS” substitution at the furin cleavage site (residues 682–685), a C-terminal T4 fibritin trimerization motif, an HRV3Cprotease cleavage site, a TwinStrepTag and an 8XHisTag in expression vector pαH was a gift from Jason McLellan (University of Texas). The plasmid was transiently transfected into Expi293 cells (Thermofisher) and Spike protein was secreted into media. Spike protein was purified using Talon resin (Takara) and concentrated to ∼1 mg/ml using Centriprep 4, aliquoted and flash frozen with liquid nitrogen.

*ACE2* purification was done using the plasmid for expressing human ACE2 as an FC-fusion (pcDNA3-sACE2(WT)-Fc(IgG1)) was a gift from Erik Procko (Addgene plasmid # 145163; http://n2t.net/addgene:145163; RRID:Addgene_145163). The plasmid was modified by introducing a TEV-cleavage site ((ENLYFQG) between the ACE2 and Fc domain. The protein was expressed and purified from Expi293 cells as described previously (1). Following purification, the Fc domain was removed by TEV-cleavage to yield the soluble monomeric ACE2 protein^29^.

*α7 nicotinic acetylcholine receptor (Alpha7)* purification was a modification of that described previously^30^. Briefly, frozen cells from 3L culture (∼45g) were thawed and resuspended in 200mls 20mM Tris (pH=7.4), 150mM NaCl, 5mM EGTA and Complete EDTA free protease inhibitor. Cells were lysed for 30 min by stirring at 4 □. Membranes were pelleted at 30,000g for 20min then resuspended in the same buffer containing 40mM n-dodecyl-β-D-maltoside (DDM; Anatrace). Following Dounce homogenization, the sample was stirred for 2 h at 4 β for protein extraction. The sample was then subjected to centrifugation (30,000g for 30min). The supernatant was applied to Strep-Tactin Superflow high-capacity resin (IBA Life Sciences) and bound receptor was washed, eluted and lipidic discs formed with the described ratios for Saposin A and and polar lipids. Detergent was removed via anion exchange using a 5ml HiTrap Q HP column (Cytiva) to bind the Alpha7-lipidic disk complex and washing with ∼20 column volumes of 20mM Tris (pH=7.4), 37.5mM NaCl and 1.25mM EGTA. The complex was eluted with a gradient to 1M NaCl. Pooled fractionss were concentrated before subjecting to size exclusion chromatography (SEC) using a Superose 6 Increase 10/300 column. Peaks fractions were use for cryo-EM grid preparation.

*α-Bungarotoxin (Btx)* (ThermoFisher Scientific, Cat. #B13422) was reconstituted in phosphate-buffered saline (PBS) to a final concentration of 1□mg/mL.

### Cryo-EM Data Acquisition and Processing

#### Common steps

All datasets used for high-resolution reconstruction were acquired on a Titan Krios G4 transmission electron microscope (Thermo Fisher Scientific) operating at 300 kV, equipped with a K3 direct electron detector and a BioContinuum™ energy filter (Gatan Inc.). Movies were preprocessed in CryoSPARC Live, applying patch motion correction and contrast transfer function (CTF) estimation using default parameters. Following data collection, micrographs were curated to remove outliers and transferred to a CryoSPARC^31^ workspace for downstream processing. Data collection parameters for all the specimens are summarized in Table 1.

Initial particle picking was performed using a blob picker with diameter ranges optimized for each dataset. Picks were curated using the “Inspect Picks” tool to exclude false positives and artifacts. Particle extraction was conducted using dataset-specific box sizes followed by Fourier cropping to reduce computational load.

Two to four rounds of 2D classification were used to discard junk particles and improve dataset quality. Initial 3D volumes were generated using ab-initio reconstruction (typically with C1 symmetry), followed by homogeneous and/or non-uniform refinement using symmetries appropriate to the target (e.g., C1, D3, O, or C5). Final refinements included local CTF refinement and reference-based motion correction. Additional advanced procedures, including symmetry expansion, focused classification, and particle subtraction, were performed for complex assemblies.

#### Apoferritin

A total of 3,760 curated micrographs were used. Initial blob picking (diameter 80–130 Å) yielded 888,579 particles, reduced to 446,824 after manual inspection. Three rounds of 2D classification resulted in 75,589 particles, from which 11 well-defined classes (27,741 particles) were selected. A single ab-initio model was generated with C1 symmetry and refined using octahedral (O) symmetry to 3.0 Å resolution. Template-based picking using these classes yielded 1,812,363 particles; after curation and re-extraction with a 480-pixel box, 86,387 unique particles remained. Subsequent refinement rounds, including local CTF and motion correction, produced a final reconstruction at 1.9 Å. Model building was performed in Phenix using PDB entry 6PXM

#### E. coli dGTPase

Preprocessing produced 1,907,816 blob-picked particles (diameter 100–150 Å), reduced to 646,655 after curation. Extraction used a 460-pixel box size, Fourier cropped to 230 pixels. One round of 2D classification yielded 141,104 particles, used for ab-initio reconstruction (C1), homogeneous refinement (C1), and non-uniform refinement (D3). Re-extracted particles (139,988 total) were refined over three rounds, resulting in a final reconstruction at 2.8 Å.

#### Spike–ACE2

After curation, 10,864 micrographs were used. Blob picking (100–230 Å) yielded 4,785,916 particles, reduced to 1,280,208 after inspection. Extracted particles (800-pixel box, cropped to 200 pixels) underwent three rounds of 2D classification, yielding 85,587 particles used to generate three ab-initio models. The best two were refined heterogeneously, identifying a class of 49,995 particles used to generate a 5.4 Å map (C1 symmetry). The resulting map served as a template for new picking (diameter 220 Å), yielding 4,402,619 particles, reduced to 2,555,824. Further processing followed similar classification and refinement steps.

#### Alpha7–Btx

A total of 25,577 curated micrographs were used. Blob picking (120–190 Å) and subsequent 2D classification generated templates for template-based picking (190 Å), yielding 10,339,021 particles. Extracted particles (540-pixel box, cropped to 180 pixels) were subjected to three rounds of 2D classification into 200 classes. 113 classes were retained (1,162,890 particles), and six ab-initio 3D classes were generated. Heterogeneous refinement identified five classes (965,331 particles) with α7-like features, used for homogeneous refinement (C5 symmetry). The consensus map was segmented, and symmetry expansion was applied. Focused 3D classification (10 classes, 6 Å filter) yielded a final set of 246,001 particles representing fully occupied complexes. Final refinement, including motion correction, resulting in a final reconstruction at 2.9 Å.

## Results and Discussion

### Blot and Plunge

We evaluated our automated cryo-EM grid preparation system by testing a range of standard and complex biological samples. To assess grid-type compatibility, we tested UltrAuFoil, Quanitfoil, HexAuFoil, and custom-fabricated gold-coated grids. In all cases, vitrification produced many grid squares with thin ice, minimal contamination, and high particle distribution, confirming consistent grid quality. These results demonstrate that the system enables reproducible high-resolution data collection across diverse sample types and supports to near-atomic structural reconstruction.

Using the blot-and-plunge method, we prepared grids under conditions detailed in Table 2. Apoferritin was reconstructed to ∼1.9 Å resolution (Fig. 3), resolving clear side chains and secondary structure elements.

**Table 2.**
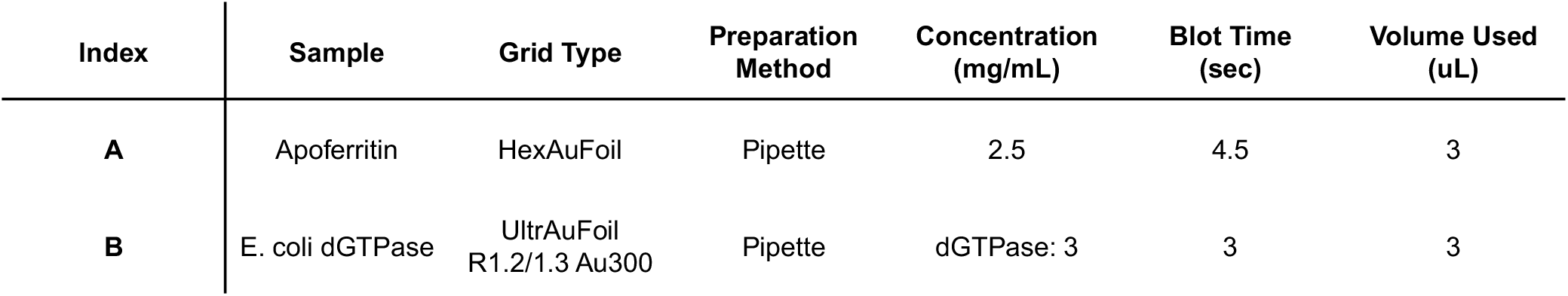
Freezing conditions for blot-and-plunge samples.

**Figure 3.**
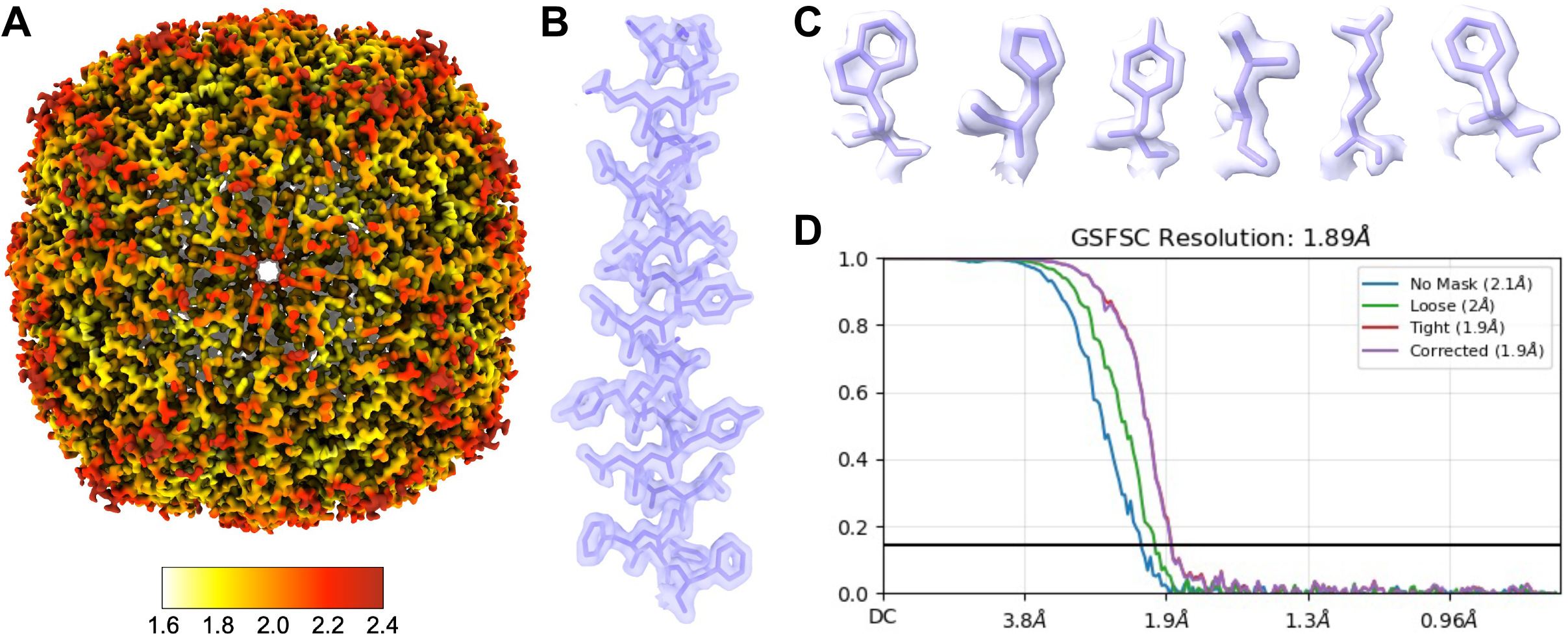
1.9 A (global estimated resolution) cryo-EM map of horse spleen apoferritin. (A) Map colored to show local resolution. (B) Representative helix (Ser9-Phe37) and (C) side chains from a single subunit with PDB: 6RJH fit to the map. (D) GSFSC curve from CryoSPARC Local Refinement showing estimated resolution calculated from two half maps.

### Spray and Blot

#### Grid Quality and Reproducibility

Spraying directly onto a dry grid often results in poor sample distribution, typically forming a thick central spot with only a narrow ring of usable squares around it, while the outer regions remain too dry to image (Fig. S3D). This unevenness severely limits the number of collectable areas and contributes to particle distribution variability.

To address this limitation, our modified approach was introduced that involves pre-wetting the grid with a small volume of buffer—typically between 500 nanoliters and 3 microliters—prior to sample spraying. The sprayed sample mixes with the buffer during the defined incubation time before plunging. Altogether, the incorporation of pre-wetting and controlled on-grid mixing strategies significantly improved sample distribution and ice uniformity supporting more consistent data collection (Fig. S3A).

On-grid mixing using minimal volumes in both spray modes enables efficient two-component reactions, offering a valuable strategy for time-resolved Cryo-EM studies. When a smaller ligand is pre-applied to the grid in molar excess and a larger biomolecule is sprayed during the blotting cycle, the interaction occurs almost immediately upon contact. In this setup, the reaction effectively outpaces diffusion, leading to higher binding efficiency. As a result, a majority of the sprayed molecules form complexes before significant dispersion can occur. This rapid and localized binding minimizes the presence of unbound particles and reduces background noise, improving particle homogeneity. This leads to more uniform particle distribution and improved consistency across grid squares, which in turn supports more reliable data collection and downstream image analysis.

The slow diffusion of the sample into the pre-applied buffer can result in concentration gradients. This is particularly problematic when working with larger macromolecules or when one of the sample components in time-resolved experiments must be present at a high molar excess to ensure interaction. In such cases, overcrowding on the grid surface can compromise in achieving good quality grid for data collection.

The integration of precision robotics with a modified on-grid mixing approach has greatly improved the reproducibility and ease of time-resolved cryo-EM experiments. This setup enables users to consistently perform rapid mixing and freezing with minimal manual intervention. However, some challenges remain—most notably in managing sample consumption and ensuring consistent spray performance. These factors depend on parameters such as sample viscosity, droplet alignment, and delivery efficiency. Ongoing optimization aims to reduce the required volume to the tens-of-nanoliters range, allowing multiple spray events from a single microliter of sample.

To maintain consistent performance across grids, the acoustic transducer surface is cleaned with ethanol after each spray. This step helps place the droplet in the center of the transducer face and avoids uneven spreading that could lead to inconsistent spraying. This simple yet critical procedure has proven highly effective: across several hundred frozen grids, we observed no evidence of cross-contamination between samples. Together, this refined workflow supports high-throughput grid preparation while preserving the integrity and reliability needed for quantitative, time-resolved studies.

Using this method, we prepared grids under conditions detailed in Table 3.

**Table 3.**
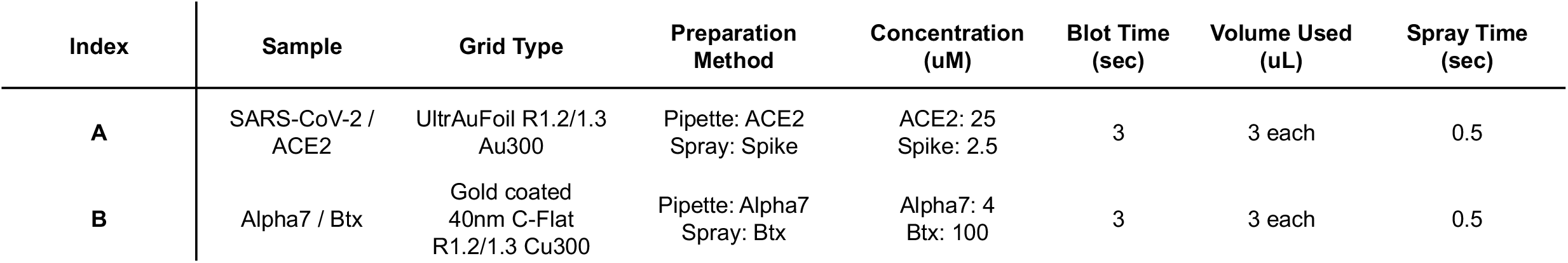
Freezing conditions for spray mixing samples.

| Index | Sample | Grid Type | Preparation Method | Concentration (uM) | Blot Time (sec) | Volume Used (uL) | Spray Time (sec) |
| --- | --- | --- | --- | --- | --- | --- | --- |
| A | SARS-CoV-2 / ACE2 | UltrAuFoil R1.2/1.3 Au300 | Pipette: ACE2<br>Spray: Spike | ACE2: 25<br>Spike: 2.5 | 3 | 3 each | 0.5 |
| B | Alpha7 / Btx | Gold coated 40nm C-Flat R1.2/1.3 Cu300 | Pipette: Alpha7<br>Spray: Btx | Alpha7: 4<br>Btx: 100 | 3 | 3 each | 0.5 |

#### Mixing SARS COV2 spike with ACE2

To evaluate the system’s performance in facilitating protein–protein complex formation, we studied the interaction between the SARS-CoV-2 spike protein and its receptor, ACE2. A 300-mesh R1.2/1.3 UltrAuFoil grid was first loaded into the robotic system. After securing the grid, 3 µL of ACE2 (2.4 mg/mL, ∼14.1 µM) was applied to the carbon side. Fast Spray mode was then initiated, triggering a 3-second blotting sequence. During the first 0.5 seconds of blotting, the spike protein (1.2 mg/mL, ∼2.7 µM trimer), pre-loaded on the back of the acoustic transducer, was sprayed onto the grid. Blotting continued for the remaining 2.5 seconds, followed immediately by plunge-freezing.

Grid screening revealed numerous areas suitable for data collection. Single-particle cryo-EM analysis yielded a reconstruction of the spike–ACE2 complex at a global resolution of 3.7□Å, based on the gold-standard FSC 0.143 criterion (Fig. 4). The dominant class showed ACE2 bound to a single receptor-binding domain (RBD) of the spike trimer, while other classes displayed two or three ACE2 molecules bound, capturing multiple receptor engagement states.

**Figure 4.**
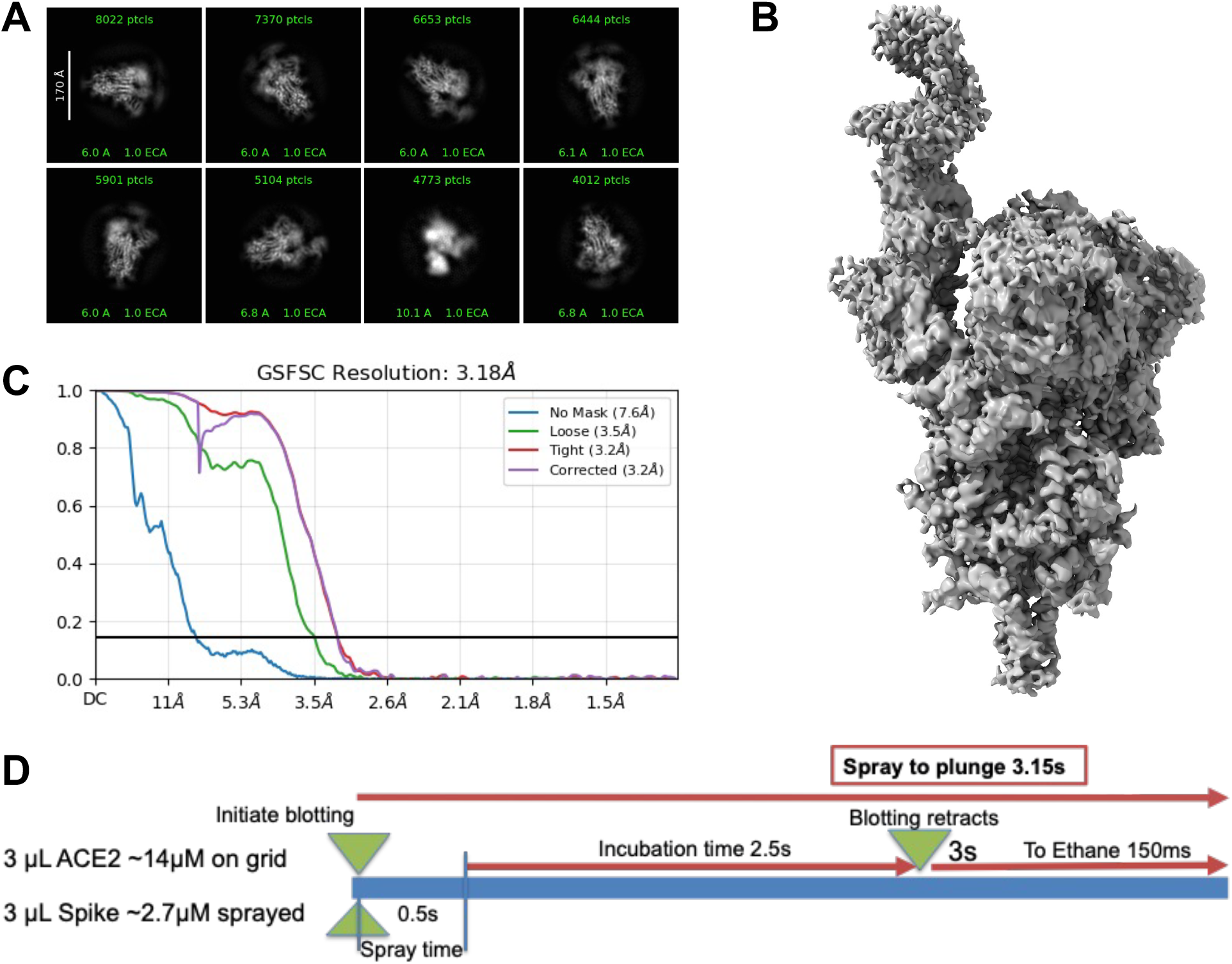
Cryo-EM results from SARS-CoV-2 Spike sprayed onto a grid containing ACE2. (A) 2D classes showing varied amounts of ACE2 bound to the spike RBD. (B) Final cryo-EM map of the spike protein with a single ACE2 bound. (C) Fourier Shell Correlation curves from two halfmaps, showing the estimated resolution of the structure shown in B. (D) Outline of experimental conditions used for freezing.

These results demonstrate that our on-grid mixing approach enables effective complex formation under low sample volumes and short incubation times, making it well-suited for transient protein–protein interaction studies.

#### Mixing Alpha7 with Btx

To investigate the mixing behavior and conformational dynamics of the α7 nicotinic acetylcholine receptor, we used the protein at a concentration of 4 µM. The sample was applied to homemade gold grids (R 1.2/1.3, 300 mesh), and α-bungarotoxin (α-Btx) was sprayed at a concentration of 100 µM for 500 ms. Grids were back blotted for 3 s prior to vitrification. Samples were screened and data were collected using a Titan Krios microscope operated at 300 kV. The recorded movies were imported into cryoSPARC for processing, including motion correction and patch-based contrast transfer function (CTF) estimation. Curated micrographs were used for particle picking and extraction, yielding a clean particle dataset. Initial ab initio reconstruction was followed by two rounds of heterogeneous refinement. Particles corresponding to 3D volumes resembling α7 nAChR were pooled and subjected to homogeneous refinement with C5 symmetry imposed. The refined map was segmented to isolate individual subunits, and signal subtraction was performed on four of the five subunits. The remaining subunit was used for symmetry expansion followed by focused 3D classification. This classification revealed multiple conformational states, which were further refined individually to obtain high-resolution structures. The final 3D reconstructions of the α7-α-Btx complex showed multiple conformations, including receptors bound to one, two, four, or five α-Btx molecules. The fully occupied receptor (bound to five α-Btx molecules) displayed the complete structure and was resolved at 2.6 Å resolution (Fig. 5). In contrast, partially occupied states primarily resolved the ectodomain, likely due to particle damage or incomplete mixing during sample preparation.

**Figure 5.**
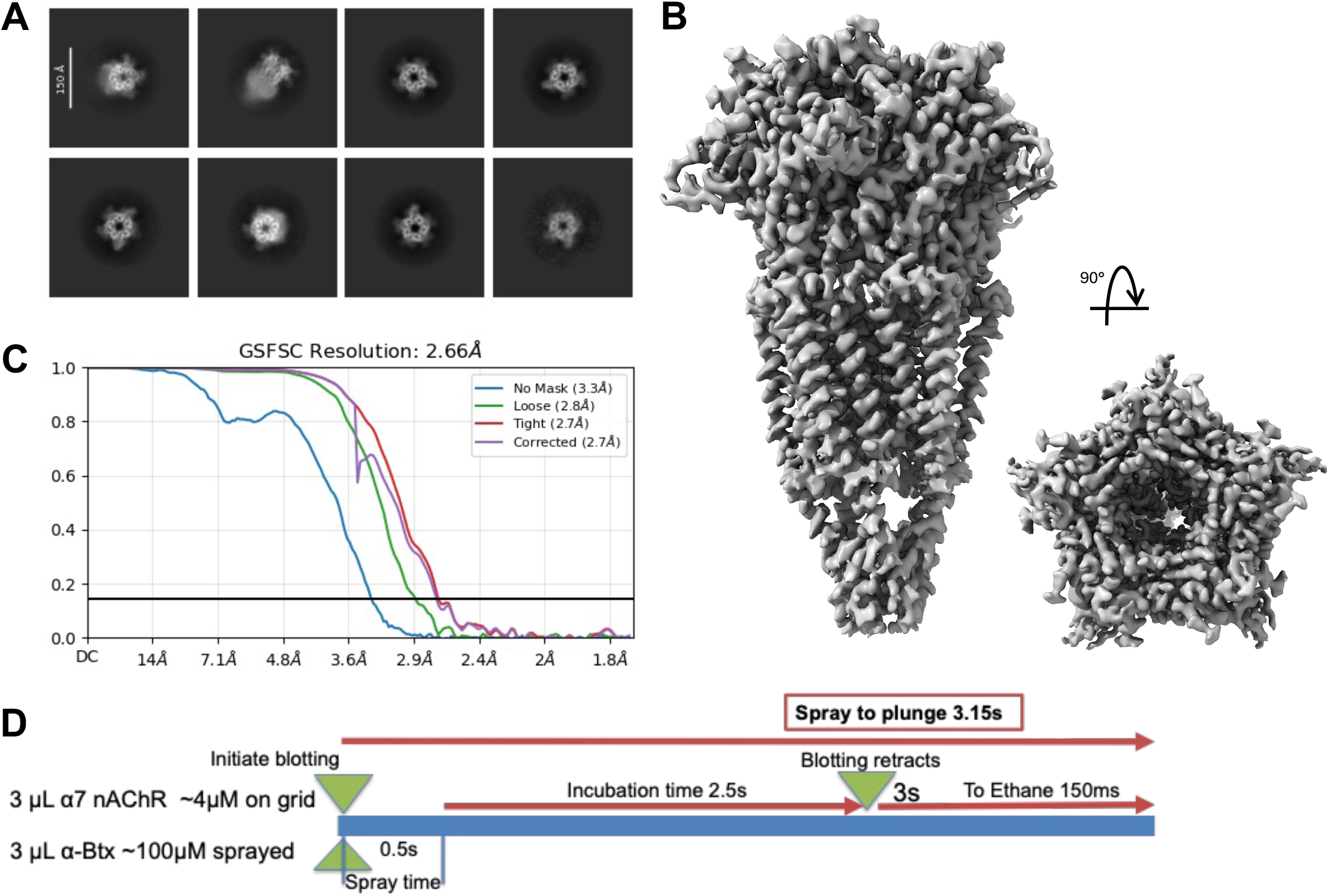
Cryo-EM results from Btx sprayed onto a grid containing Alpha7. (A) 2D classes showing varied amounts of Btx bound to Alpha7. (B) Final cryo-EM map of five Btx found to a single Alpha7. (C) Fourier Shell Correlation curves from two half-maps, showing the estimated resolution of the structure shown in B. (D) Outline of experimental conditions used for freezing.

## Conclusion

Cryo-EM continues to rapidly evolve as a powerful structural biology technique, yet current grid preparation methods remain restrictive and inflexible. Although recent advancements have introduced improvements, many commercial solutions remain limited in adaptability, quickly becoming obsolete as the demands of cryo-EM research expand and diversify. To address these challenges, we developed VitriFlex—a modular, open-source cryo-EM grid preparation system designed to not only fulfill current research needs but adapt flexibly to future developments.

We validated VitriFlex’s efficacy by achieving near-atomic resolution reconstructions of multiple proteins, including apoferritin and dGTPase. The system’s versatility was further demonstrated through various mixing experiments including spike protein with ACE2 and Alpha7 with Btx.

VitriFlex already incorporates advancements, including automated grid transfers and digital logging of freezing parameters, which reduce user variability and enhance reproducibility. Future work will focus on optimizing sample mixing both prior to and directly on the grid, along with exploring active liquid-thinning methods to further decrease plunge times while maintaining a sufficient number of collectible grid areas. Integration with web-based grid screening software, such as SmartScope, is also planned, aiming to streamline grid evaluation and simplify freezing condition optimization, making the entire workflow more accessible and reliable.

VitriFlex is offered as an open-source platform, promoting community engagement to continuously expand its capabilities through the development of additional modules and functionalities. With substantial cost savings and minimal technical expertise required for setup and operation, VitriFlex significantly lowers the barrier to entry for cryo-EM research. By democratizing advanced sample preparation methods, we aim to broaden access, support a wider array of innovative experiments, and continue pushing the boundaries of structural biology research.

## Acknowledgements

This research was supported by the Intramural Research Program of the National Institutes of Health (NIH); US National Institutes of Environmental Health Sciences (ZIC ES103326 to M.J.B.). The contributions of the NIH author(s) were made as part of their official duties as NIH federal employees, are in compliance with agency policy requirements, and are considered Works of the United States Government. However, the findings and conclusions presented in this paper are those of the author(s) and do not necessarily reflect the views of the NIH or the U.S. Department of Health and Human Services.

We are grateful for the technical support from Kaichun Yang from Dr. Tony Huang’s lab in Duke University’s Pratt School of Engineering, and Furkan Tasdelen from Dr. Mehmet Sen’s lab in the University of Houston’s School of Natural Science and Mathematics. We also want to thank Dave Shock from Dr. Roel Schaaper’s group for preparation of dGTPase samples.

## List of Abbreviations

dGTPase: *Escherichia coli* dGTP Triphosphohydrolase
Spike: SARS-CoV-2 S-protein
ACE2: Angiotensin-converting enzyme 2
Btx: Alpha-Bungarotoxin
RBD: Receptor-binding domain

## References

1. Nakane, T. et al. Single-particle cryo-EM at atomic resolution. NATURE 587, 152–156 (2020).

2. Bai, X., McMullan, G. & Scheres, S. H. W. How cryo-EM is revolutionizing structural biology. Trends Biochem Sci 40, 49–57 (2015).

3. Fernandez-Leiro, R. & Scheres, S. H. W. Unravelling biological macromolecules with cryo-electron microscopy. Nature 537, 339–346 (2016).

4. Kühlbrandt, W. The Resolution Revolution. Science 343, 1443–1444 (2014).

5. Dubochet, J. et al. Cryo-electron microscopy of vitrified specimens. Q Rev Biophys 21, 129–228 (1988).

6. Ravelli, R. B. G. et al. Cryo-EM structures from sub-nl volumes using pin-printing and jet vitrification. Nat Commun 11, 2563 (2020).

7. Wei, H. et al. Optimizing ‘self-wicking’ nanowire grids. J Struct Biol 202, 170–174 (2018).

8. Gilkey, J. C. & Staehelin, L. A. Advances in ultrarapid freezing for the preservation of cellular ultrastructure. Journal of Electron Microscopy Technique 3, 177–210 (1986).

9. Henderikx, R. J. M. et al. VitroJet: new features and case studies. Acta Crystallogr D Struct Biol 80, 232–246 (2024).

10. Levitz, T. S. et al. Approaches to Using the Chameleon: Robust, Automated, Fast-Plunge cryoEM Specimen Preparation. Front Mol Biosci 9, 903148 (2022).

11. Razinkov, I. et al. A new method for vitrifying samples for cryoEM. Journal of Structural Biology 195, 190–198 (2016).

12. Dandey, V. P. et al. Spotiton: New features and applications. J Struct Biol 202, 161–169 (2018).

13. Dandey, V. P. et al. Time-resolved cryo-EM using Spotiton. Nat Methods 17, 897–900 (2020).

14. Bhattacharjee, S. et al. Time resolution in cryo-EM using a PDMS-based microfluidic chip assembly and its application to the study of HflX-mediated ribosome recycling. Cell 187, 782-796.e23 (2024).

15. Feng, X. & Frank, J. Time-resolved cryo-EM (TRCEM) sample preparation using a PDMS-based microfluidic chip assembly. 2024.12.08.627437 Preprint at 10.1101/2024.12.08.627437 (2024).

16. Feng, X. et al. A fast and effective microfluidic spraying-plunging method for high-resolution single-particle cryo-EM. Structure 25, 663–670 (2017).

17. Mäeots, M.-E. et al. Modular microfluidics enables kinetic insight from time-resolved cryo-EM. Nat Commun 11, 3465 (2020).

18. Torino, S., Dhurandhar, M., Stroobants, A., Claessens, R. & Efremov, R. G. Time-resolved cryo-EM using a combination of droplet microfluidics with on-demand jetting. Nat Methods 20, 1400–1408 (2023).

19. Tan, Y. Z. & Rubinstein, J. L. Through-grid wicking enables high-speed cryoEM specimen preparation. Acta Crystallogr D Struct Biol 76, 1092–1103 (2020).

20. Rubinstein, J. L. et al. Shake-it-off: a simple ultrasonic cryo-EM specimen-preparation device. Acta Crystallogr D Struct Biol 75, 1063–1070 (2019).

21. Liu, N. & Wang, H.-W. Better Cryo-EM Specimen Preparation: How to Deal with the Air-Water Interface? J Mol Biol 435, 167926 (2023).

22. Noble, A. J. et al. Reducing effects of particle adsorption to the air-water interface in cryo-EM. Nat Methods 15, 793–795 (2018).

23. Zheng, L. et al. Self-assembled superstructure alleviates air-water interface effect in cryo-EM. Nat Commun 15, 7300 (2024).

24. Lu, Y. et al. Functionalized graphene grids with various charges for single-particle cryo-EM. Nat Commun 13, 6718 (2022).

25. Xu, J. et al. Graphene sandwich–based biological specimen preparation for cryo-EM analysis. Proceedings of the National Academy of Sciences 121, e2309384121 (2024).

26. Yang, Z. et al. Electrospray-assisted cryo-EM sample preparation to mitigate interfacial effects. Nat Methods 21, 1023–1032 (2024).

27. Fan, X. et al. Single particle cryo-EM reconstruction of 52 kDa streptavidin at 3.2 Angstrom resolution. Nat Commun 10, 2386 (2019).

28. Mechanism by which T7 bacteriophage protein Gp1.2 inhibits Escherichia coli dGTPase | PNAS. https://www.pnas.org/doi/10.1073/pnas.2123092119.

29. Engineering human ACE2 to optimize binding to the spike protein of SARS coronavirus 2 | Science. https://www.science.org/doi/10.1126/science.abc0870.

30. Noviello, C. M. et al. Structure and gating mechanism of the α7 nicotinic acetylcholine receptor. Cell 184, 2121-2134.e13 (2021).

31. Punjani, A., Rubinstein, J. L., Fleet, D. J. & Brubaker, M. A. cryoSPARC: algorithms for rapid unsupervised cryo-EM structure determination. Nat Methods 14, 290–296 (2017).

