## Supplemental Figures for "VitriFlex: An Open-Source, Modular, and Customizable Robotic Platform for Cryo-EM Grid Preparation"

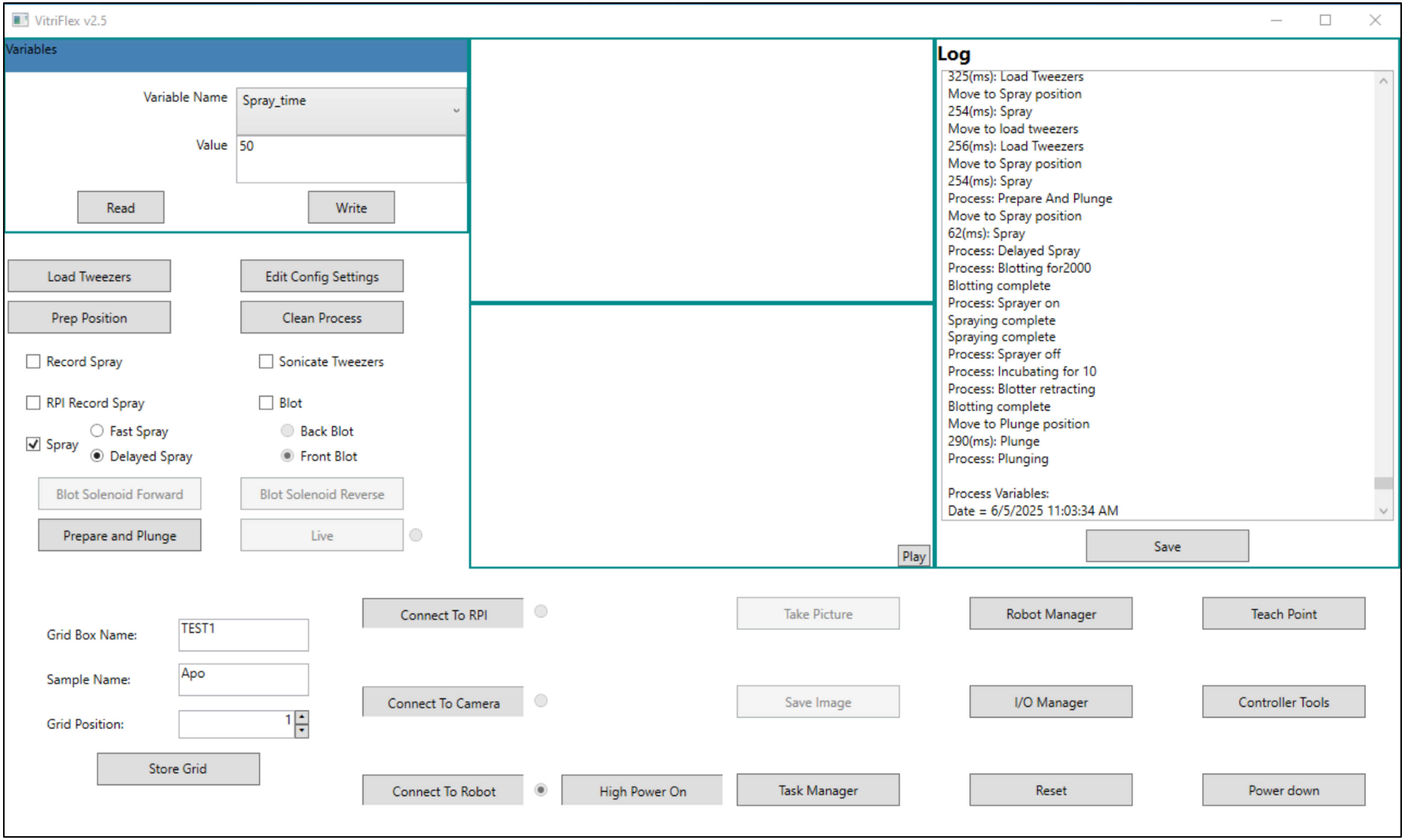


**S1. VitriFlex GUI**

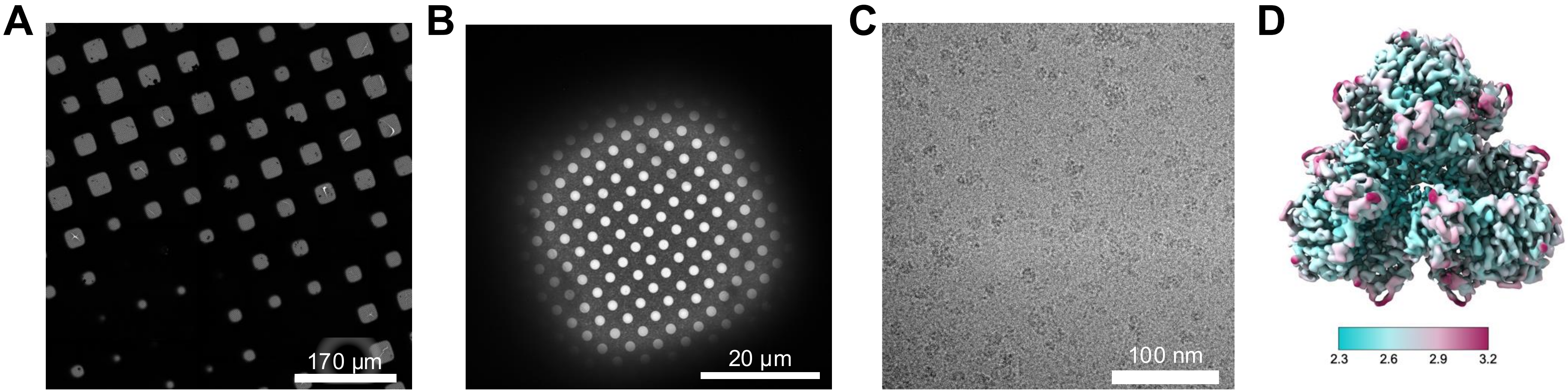


**S2. Multi-scale visualization of cryo-EM data collection for *E. coli* dGTPase.** (A) Partial grid atlas, (B) grid square, (C) representative micrograph, and (D) final 3D reconstruction. The structure was determined at an estimated global resolution of 2.8 Å, with local resolution shown in angstroms.

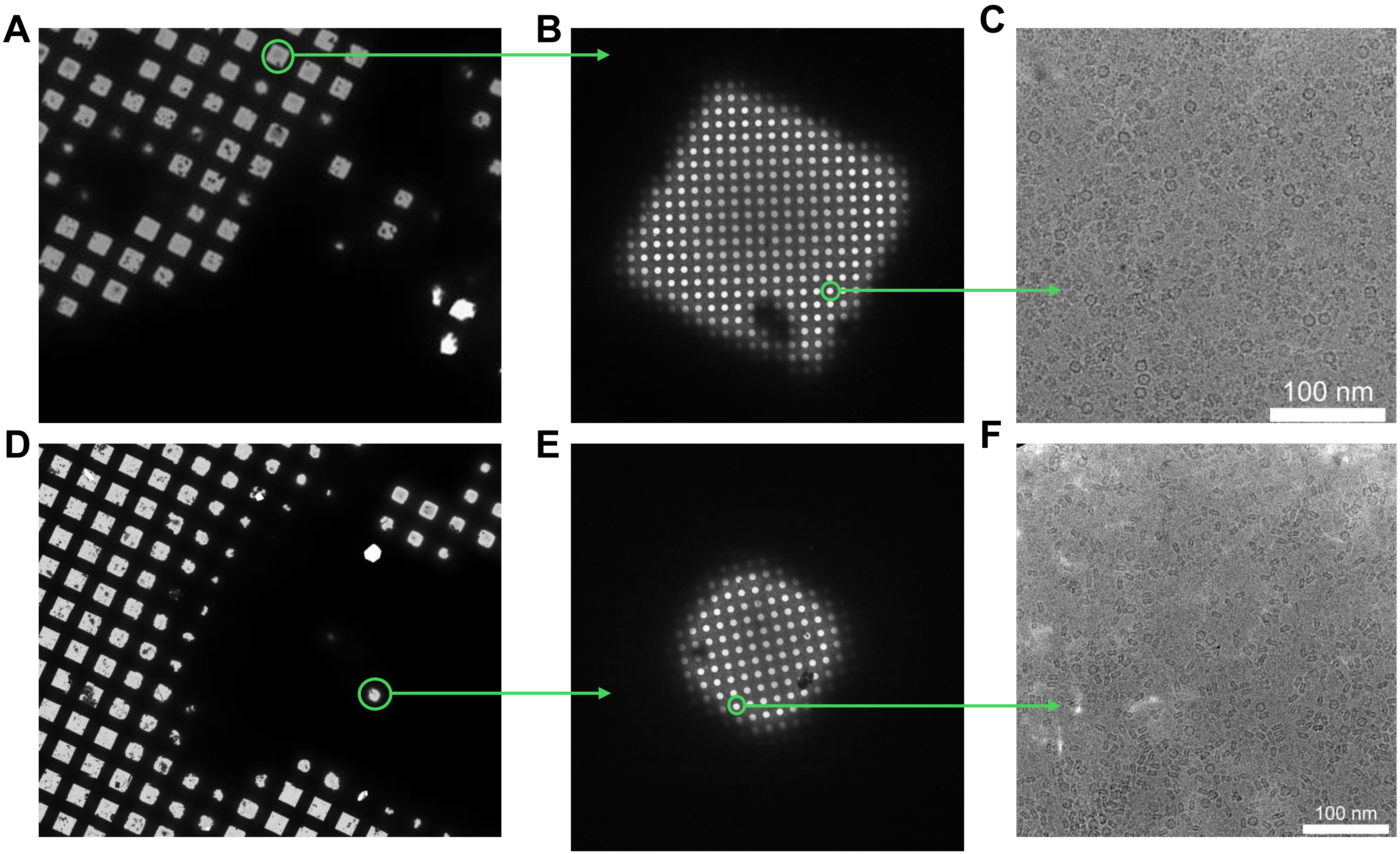


**S3. Partial atlas and high-mag cryo-EM images of spray mixing grids prepared with (A) and without (D) pre-wetting.** The grid shown in A-C was prepared with 50 ms spray of dGTPase on a grid with 1 uL apoferritin and an incubation of 10ms before plunging into liquid ethane. The grid shown in D-F was prepared by placing dGTPase and apoferritin on the sprayer, spraying for 30 ms and plunging into liquid ethane immediately following the spray.

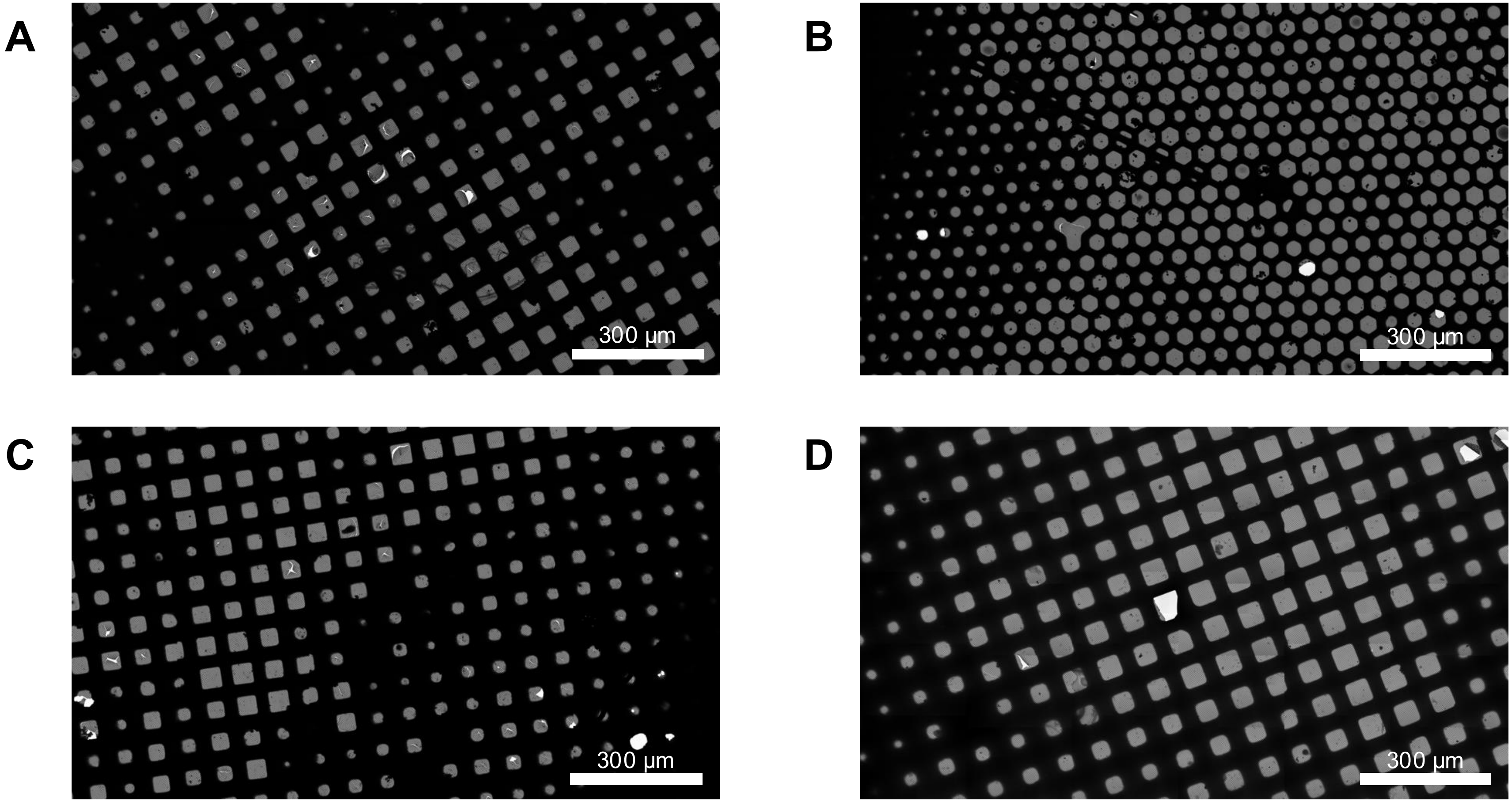


**S4. Partial atlases of cryo-EM grids used for data collection. (A),** dGTPase frozen on UltrauFoil and **(B),** apoferritin frozen on HexAuFoil, both using manual application. **(C),** Spike-ACE2 and **(D)** Alpha7-Btx, both frozen on UltrAuFoil using spray application.

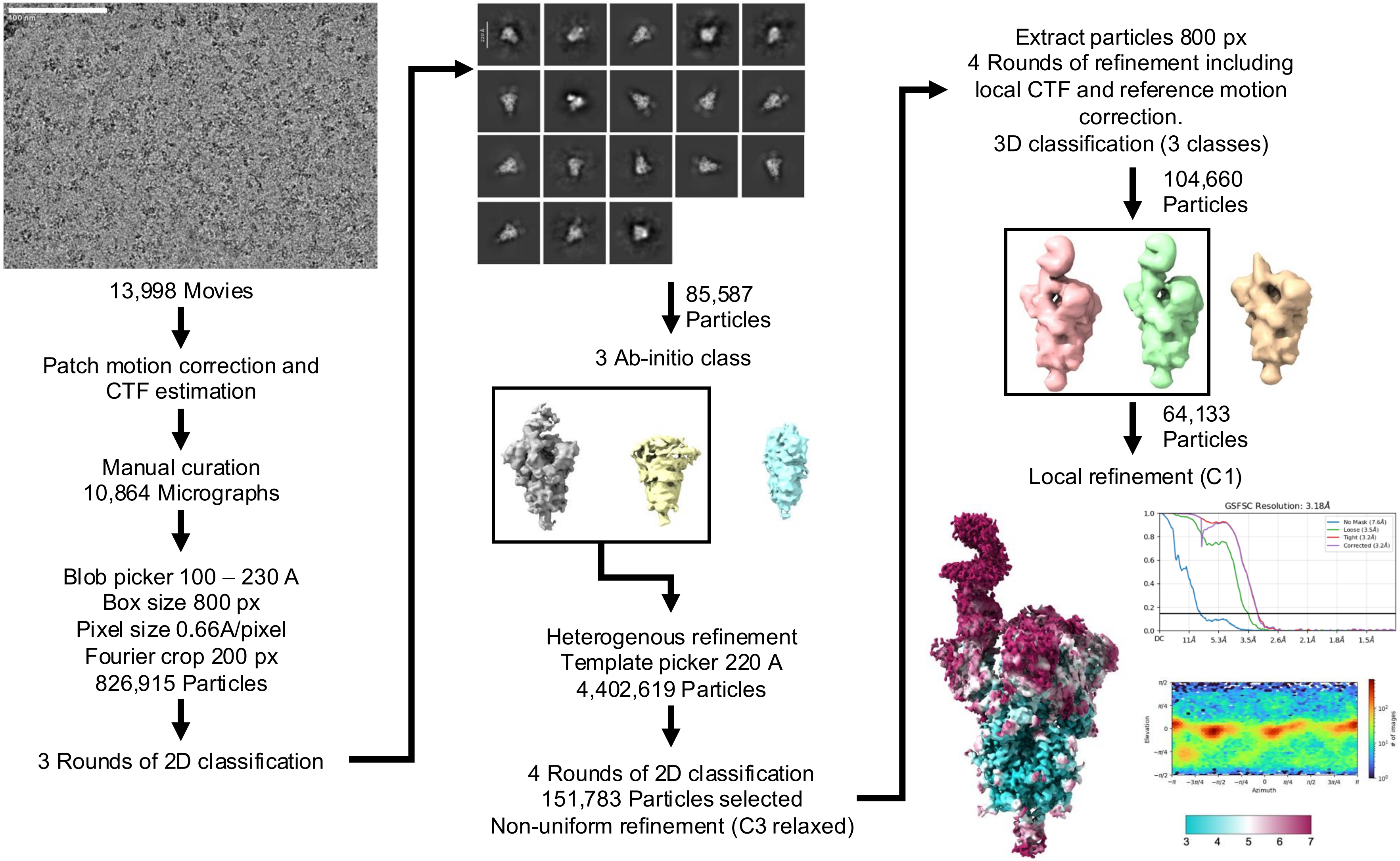


**S5. Cryo-EM processing workflow for SARS-CoV-2 Spike and ACE2 mixing experiment.**

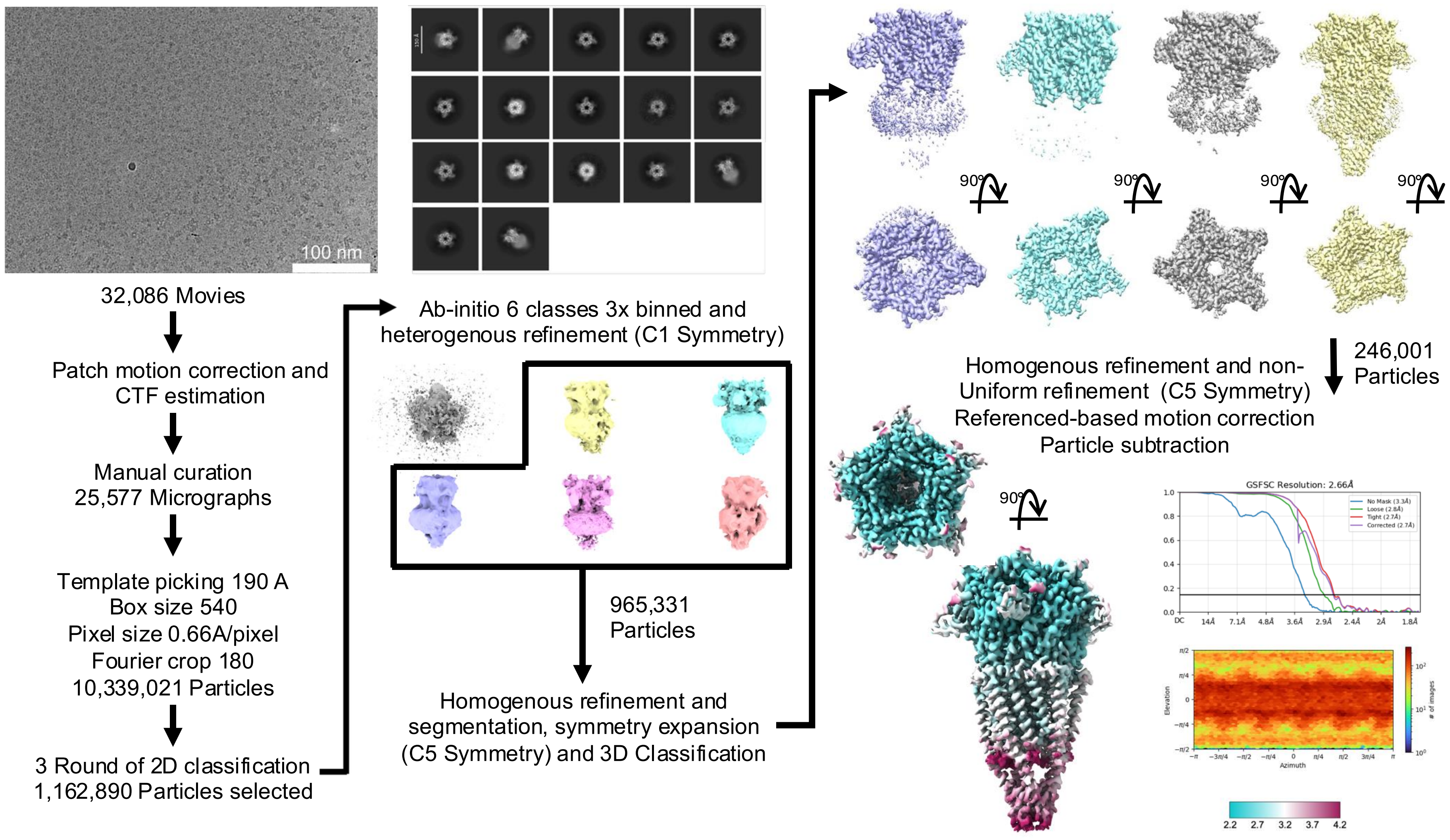


**S6. Cryo-EM processing workflow for Alpha7 and Btx mixing experiment.**
